# Epithelial-to-mesenchymal transition activates Bcat1 expression to promote recurrent tumor growth

**DOI:** 10.1101/2020.12.08.416479

**Authors:** Douglas B. Fox, James V. Alvarez

## Abstract

Branched-chain amino acid metabolism has emerged as a crucial regulator of tumor proliferation. In particular, the branched-chain amino acid transaminase 1 (BCAT1), which catalyzes the conversion of branched-chain amino acids to α-keto acids in the cytoplasm, has been identified as a promising therapeutic target for several different cancer types, including gliomas and leukemias. BCAT1 expression has also been associated with resistance to anti-estrogen therapy in breast cancer, suggesting a role in therapeutic resistance and tumor progression. Yet a functional role for BCAT1 in promoting breast cancer recurrence has not been described. Further, BCAT1 expression is restricted to just a few specialized tissues, such as the testes and central nervous system, and the mechanism by which it becomes transcriptionally activated in tumors that arise from tissues that do not typically express BCAT1 is unclear. Here, we report that Bcat1 is upregulated in recurrent breast tumors to promote recurrent tumor growth. We find that Bcat1 is upregulated by the epithelial-to-mesenchymal transition (EMT) in mammary epithelial cells and breast cancer cells. Finally, we analyze human tumor datasets to show that BCAT1 expression is strongly correlated with epithelial-to-mesenchymal gene signatures, suggesting that EMT is the predominant regulator of BCAT1 expression in cancer. Because EMT is associated with therapy resistance, metastasis, and tumor recurrence, these findings suggest a broad clinical potential for BCAT1 inhibition.

## Introduction

Leucine, isoleucine, and valine are categorized as branched-chain amino acids (BCAAs) because of their branched hydrocarbon functional groups. BCAAs are essential amino acids that are required for protein synthesis, but they can also be catabolized to capture energy from the hydrocarbon functional groups. The first enzymatic step of BCAA catabolism is a reversible transamination, where the amino group of a BCAA and the oxygen from the keto group of alpha-ketoglutarate (αKG) are exchanged [1]. This reaction forms glutamate and the branched-chain ketoacid (BCKA) derivative. BCKA oxidation occurs in the mitochondria and begins with irreversible dehydrogenation by branched-chain ketoacid dehydrogenase [2]. The ultimate fate of dehydrogenated BCKAs is to form acetyl-coenzyme A and enter the citric acid cycle to generate ATP [1]. Thus, BCAA metabolism by this catabolic pathway is a mechanism for using carbon-rich BCAAs to provide energy.

There are two BCAA transaminases (BCATs) that catalyze the initial reversible transamination reaction. BCAT2, encoded by the *Bcat2* gene, is localized in the mitochondria and is ubiquitously expressed. SLC25A44 is a mitochondrial BCAA transporter that is necessary for mitochondrial transamination of BCAAs by BCAT2 [3]. BCAT1, encoded by the *Bcat1* gene, is localized to the cytosol and has a more restricted expression pattern, with expression in the brain, ovary, and kidney [4]. However, the role of the different subcellular localization of BCAT1 and BCAT2 is not fully understood.

BCAAs have recently emerged as important nutrients for tumor cell proliferation [2], and diverse functional roles for BCAAs in tumor metabolism have been reported. Dietary BCAAs are required to sustain the high demand for protein synthesis characteristic of tumor cells [1], but they can also be used as a nitrogen source nucleotide synthesis in lung cancer cells [5]. BCAT enzymes can also provide glutamate pools necessary for glutathione synthesis, contributing to maintenance of redox homeostasis [6]. Additionally, BCAT1 activity in leukemia cells has been shown to restrict αKG levels and thereby inhibit αKG-dependent enzymes, including the TET family of DNA demethylases and prolyl hydroxylases that regulate the degradation of hypoxia inducible factors [7]. However, it has also been reported that the reverse reaction, in which BCAT1 catalyzes the transamination of plasma BCKAs to generate BCAAs, maintains nutrient sensing through mTORC1 to sustain proliferative signaling in leukemia cells [8]. Together, these recent studies demonstrate that BCAT1 represents a promising therapeutic target for tumors in which it is expressed. However, a systematic understanding of what tumor types express and are dependent upon BCAT1 activity is lacking. In non-disease states, BCAT1 expression is limited to the brain, testes, and ovaries [9], and although BCAT1 expression been shown to be modulated by c-MYC [10–12] and HIF-1α [13], it remains unknown how BCAT1 is transcriptionally activated in tumor cells, and the molecular mechanisms that underlie BCAT1 dependencies in diverse tumor types is unclear.

BCAT1 was shown to be upregulated in circulating tumor cells isolated from patients with hepatocellular carcinoma [14] and in breast cancer cells that have acquired resistance to anti-estrogen therapy [15], suggesting that BCAT1 upregulation promotes tumor progression and therapeutic resistance. However, a functional role for BCAT1 in promoting the growth of therapy-resistant recurrent tumors has not been established. Here, using a genetically engineered mouse model of breast cancer recurrence, we find that Bcat1 is dramatically upregulated during tumor recurrence and identify a functional role for BCAT1 in recurrent breast cancer. Furthermore, using both mechanistic studies in mouse tumor cells and expression data from a pan-cancer analysis of human tumors, we identify EMT as a common mechanism for BCAT1 activation in tumor cells.

## Results

### Bcat1 is upregulated in recurrent breast cancer

We previously identified an antioxidant metabolic pathway that promotes breast cancer recurrence [16]. To determine if other metabolic pathways were altered during tumor recurrence, we used a transgenic mouse model of Her2-driven breast cancer [17–19]. In this model, doxycycline administration to bitransgenic MMTV-rtTA;TetO-Her2 (MTB;TAN) mice induces mammary-specific expression of Her2 and formation of invasive mammary adenocarcinomas. Removal of doxycycline after tumor formation causes Her2 downregulation, inducing tumor regression, but a small population of cells survives. These cells persist in a dormant state [4, 20, 21] before eventually re-initiating proliferation to form recurrent tumors. To assess metabolic pathways associated with tumor recurrence, we performed gene expression analysis of two primary and two recurrent tumor cell lines. The majority of genes remained mostly unchanged between primary and recurrent tumor cell lines, however 2,414 genes (of 11,313 detected genes) were substantially altered (fold change > 2, p < 0.05) (**Fig. 1a**), demonstrating that there are specific gene expression changes associated with tumor recurrence in this model.

**Fig. 1.**
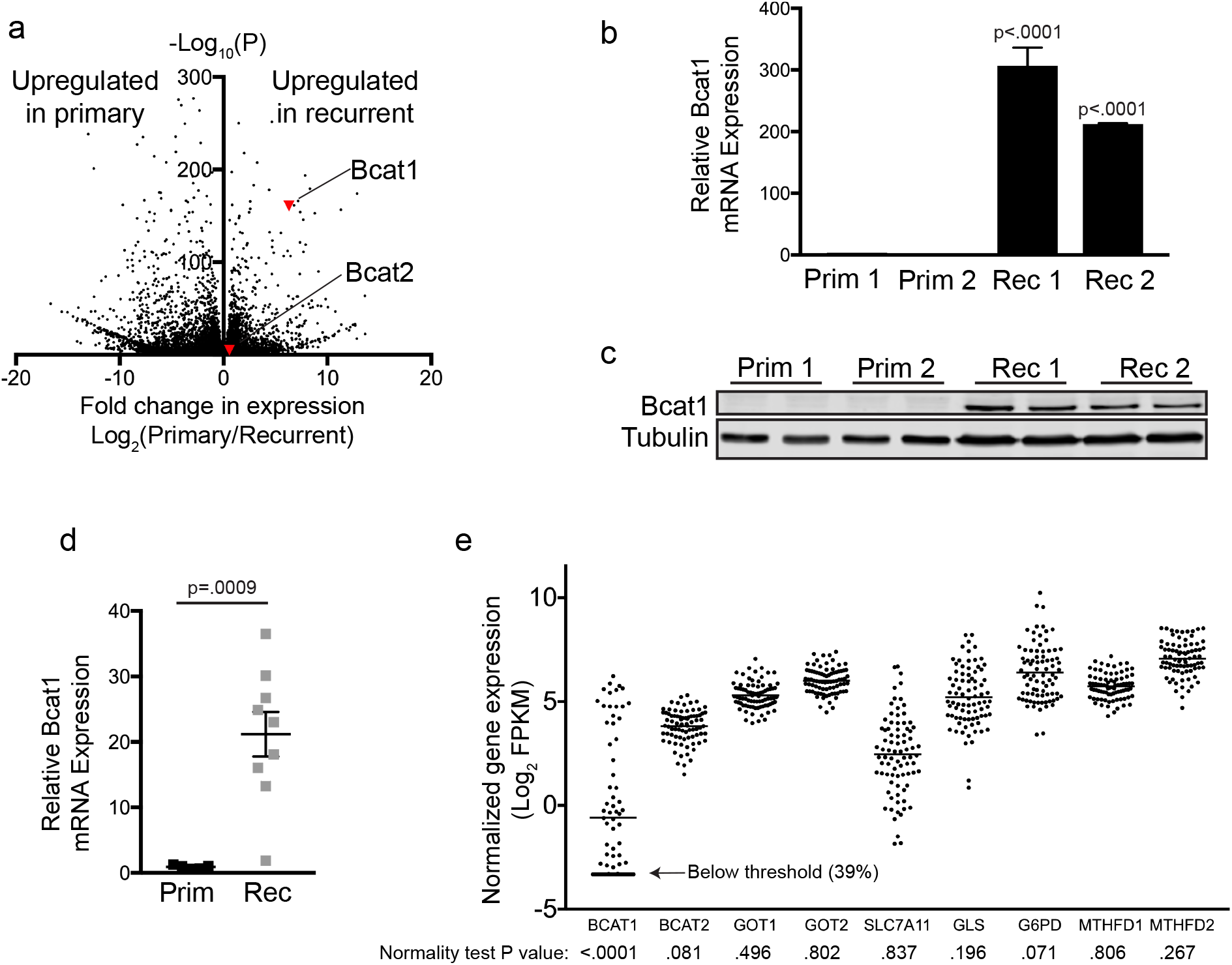
Bcat1 is upregulated in recurrent breast cancer. **a**, Volcano plot showing gene expression changes between primary (n=2) and recurrent (n=2) tumor cell lines. Black dots represent a single gene, and Bcat1 and Bcat2 are labeled with red triangles. **b**, qRT–PCR analysis of Bcat1 expression in primary (n=2) and recurrent (n=2) tumor cell lines. Mean ± SEM for n=3 biologically independent replicates. Significance was determined by one-way ANOVA (Tukey’s multiple comparisons test). **c**, Western blot for BCAT1 in two independent primary and recurrent cell lines. **d**, qRT–PCR analysis of Bcat1 expression for primary (n=5) and recurrent (n=8) MTB;TAN tumors. Data are represented as mean ± SEM. Significance was determined by Student’s t-test. **e**, Plot showing expression levels for a panel of metabolism genes from a panel of 86 breast cancer cell lines. Each dot represents the expression value for a single cell line and the black bar represents the mean. Distribution normality was tested using the Shapiro-Wilk normality test, where p<0.05 indicates a significant deviation from a normal distribution.

Strikingly, Bcat1 expression was ranked among the top of all genes upregulated in recurrent tumor cells (**Fig. 1a**). To validate the transcriptomic analysis, we performed qRT-PCR using independent RNA samples. Bcat1 expression was greater than 200-fold higher in recurrent tumor cell lines (**Fig. 1b**), and western blotting showed that BCAT1 protein was highly expressed in recurrent cells, but undetectable in primary tumor cells (**Fig. 1c**). Additionally, qRT-PCR analysis of primary and recurrent tumor samples was used to confirm that Bcat1 is upregulated in recurrent tumors *in vivo* (**Fig. 1d**). Together, these results suggest that Bcat1 expression is induced by tumor recurrence. Intrigued by the binary expression pattern of Bcat1 in MTB;TAN tumor cells, we examined the pattern of Bcat1 expression in human breast cancer cells using RNA-seq data from a panel of 86 human breast cancer cell lines [22]. For reference, we compared BCAT1 expression to the expression of other transaminases and metabolic enzymes commonly expressed in cancer cells. Whereas the expression of most metabolic enzymes followed a normal distribution, BCAT1 had a bimodal expression pattern and was not normally distributed (**Fig 1e;** p<0.0001 Shapiro-Wilk normality test). Further, Bcat1 expression was below the detection threshold in 39% of cell lines (**Fig. 1e**). This analysis demonstrates that, unlike other metabolic enzymes, BCAT1 is not expressed or expressed at very low levels in a large subset of breast cancer cell lines.

### Bcat1 promotes recurrent tumor growth

We next examined the functional importance of Bcat1 for tumor recurrence. To do this, we transduced recurrent tumor cells to express Cas9 and a guide RNA (gRNA) targeting either Bcat1- or GFP. Western blotting confirmed that BCAT1 protein expression was ablated by both gRNAs targeting Bcat1 (**Fig. 2a**). Bcat1 knockout had no effect on the proliferation rate of recurrent tumor cells *in vitro* (**Fig. 2b**). However, when recurrent tumor cells were orthotopically implanted into the mammary fat pad of recipient mice, tumor growth was impaired in Bcat1-knockout tumors (**Fig. 2c,d**). These results demonstrate that Bcat1 expression is activated during tumor recurrence and is functionally important for the growth of recurrent tumors *in vivo*. Interestingly, these results agree with a study showing knockout of BCAT1 and BCAT2 in non-small cell lung cancer cells affects tumor growth *in vivo* but not tumor cell proliferation *in vitro* [5].

**Fig. 2.**
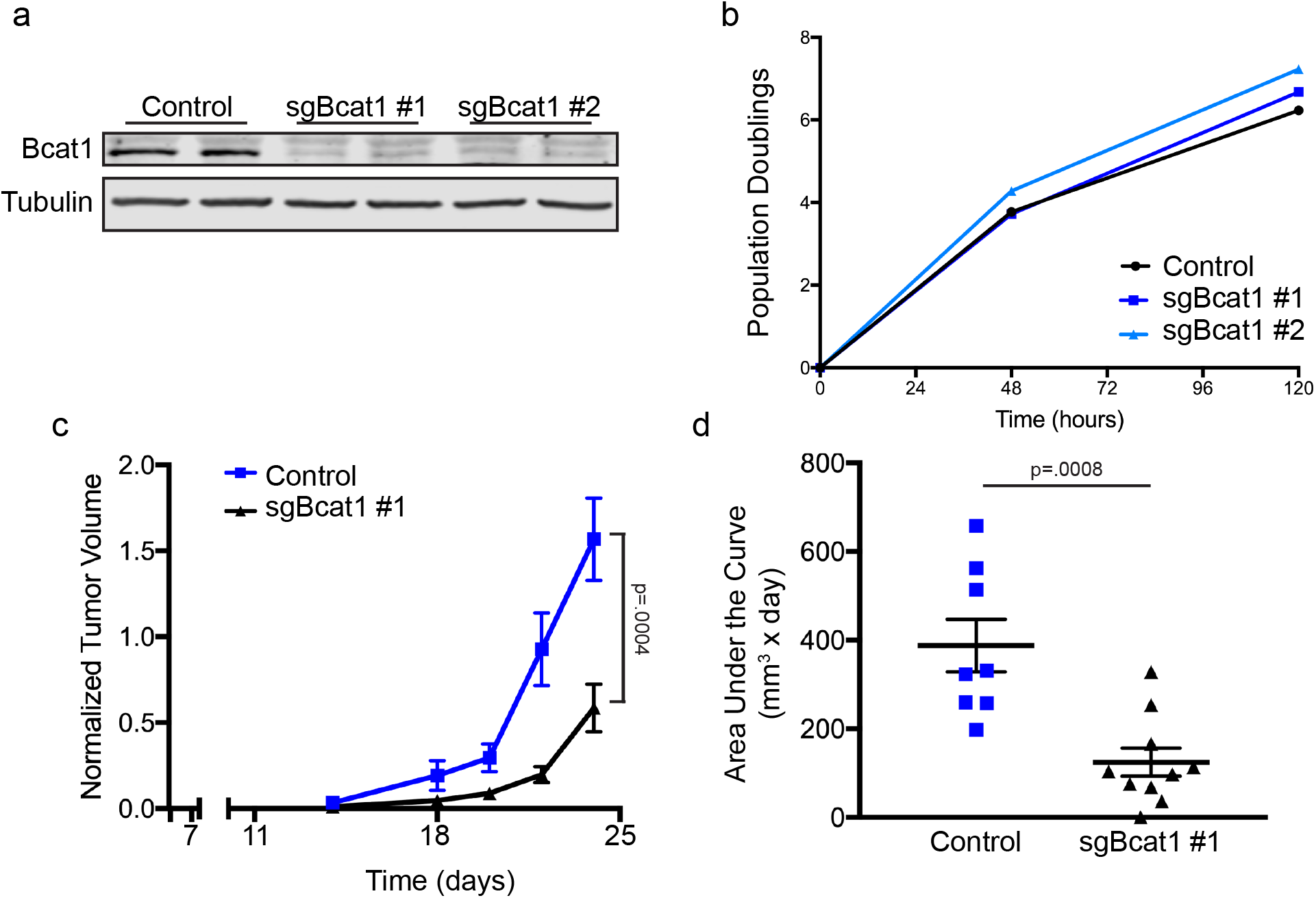
Bcat1 promotes recurrent tumor growth. **a**, Western blot for BCAT1 in control (sgGFP) and BCAT1-knockout (sgBcat1) recurrent tumor cells. **b**, Population doublings in control (sgGFP) and BCAT1-knockout (sgBcat1) recurrent tumor cells. n=2 biological replicates for each time point. Data are representative of two independent experiments **c**,**d**, Tumor growth curves (**c**) and area under the curve analysis (**d**) for tumors generated from control (sgGFP, n=8 tumors) and Bcat1-knockout (shBcat1, n=10 tumors) recurrent tumor cells. Significance was determined by Student’s t-test using the mean tumor volumes (**c**) and the mean area under the curve values (**d**) at experiment endpoint (n=8 control tumors, n=10 sgBcat1 tumors). Data are representative of a single experiment.

### Bcat1 expression is activated by TGF-β induced epithelial-to-mesenchymal transition

Considering the dramatic difference in Bcat1 gene expression between primary and recurrent tumors, we next investigated how Bcat1 is transcriptionally regulated. We first used chromatin immunoprecipitation sequencing (ChIP-seq) to analyze the occupancy of RNA Polymerase II (RNA Pol II) and the relative abundance of repressive and activating post-translation histone modifications at the Bcat1 genetic locus. In agreement with qRT-PCR analysis (**Fig. 1b**), ChIP-seq revealed high RNA Pol II occupancy at the Bcat1 promoter in recurrent tumor cells (**Extended Data Fig. 1a).** In contrast, no RNA Pol II was detected at the Bcat1 locus in primary tumor cells (**Extended Data Fig. 1a**). Similarly, the activating histone modifications histone H3 lysine 9 acetylation (H3K9Ac) and histone H3 lysine trimethylation (H3K4me3) were enriched at the Bcat1 promoter in recurrent cells, but were scarcely detected in primary cells (**Extended Data Figs. 1b,c**). Conversely, the repressive histone modification H3 lysine 27 trimethylation (H3K27me3) was more abundant throughout the Bcat1 gene in primary cells (**Extended Data Figs. 1d**). Together, these results demonstrate that elevated Bcat1 expression in recurrent tumor cells is associated with epigenetic remodeling at the Bcat1 locus, consistent with an “on/off” pattern of Bcat1 expression.

Primary tumors arising in MTB;TAN mice exhibit an epithelial morphology characteristic of primary breast adenocarcinomas [18], and tumor cells cultured from these tumors maintain this morphology [23]. In contrast, recurrent tumors in MTB;TAN mice, and cell lines cultured from these tumors, display morphological changes characteristic of epithelial-to-mesenchymal transition (EMT) [17, 18]. Gene set enrichment analysis showed that the Hallmark gene signature “epithelial-to-mesenchymal transition” is markedly enriched in recurrent tumor cells (**Fig. 3a**). EMT often involves dramatic epigenetic remodeling to induce expression of EMT effector genes, such as Snail, Twist, and Cdh2, that are silenced in epithelial cells [24]. Further, BCAT1 can be regulated by the EMT transcription factor, SMAD5, in fibroblasts [25]. We therefore considered that Bcat1 upregulation in recurrent tumor cells might be a consequence of EMT. To test this, we used TGF-β to induce EMT in two primary tumor cell lines. We observed transformation to a spindle-shaped morphology in most cells after 10 days of TGF-β treatment, while vehicle treated cells maintained epithelial morphology (**Fig. 3b,c**). EMT was confirmed in these cells using qRT-PCR analysis of canonical EMT markers. TGF-β treatment induced silencing of the epithelial marker Cdh1 in both primary tumor cell lines (**Fig. 3d,e**). Expression of mesenchymal markers Twist1 and Snail1 was upregulated by TGF-β in one primary tumor cell line, and the mesenchymal markers Snail1 and Vimentin were induced in the other (**Fig. 3d,e**), demonstrating that these cells had undergone transcriptional changes characteristic of EMT. Importantly, Bcat1 expression was induced 285- and 18-fold, respectively, in TGF-β treated primary tumor cells (**Fig. 3f,g**).

**Fig. 3.**
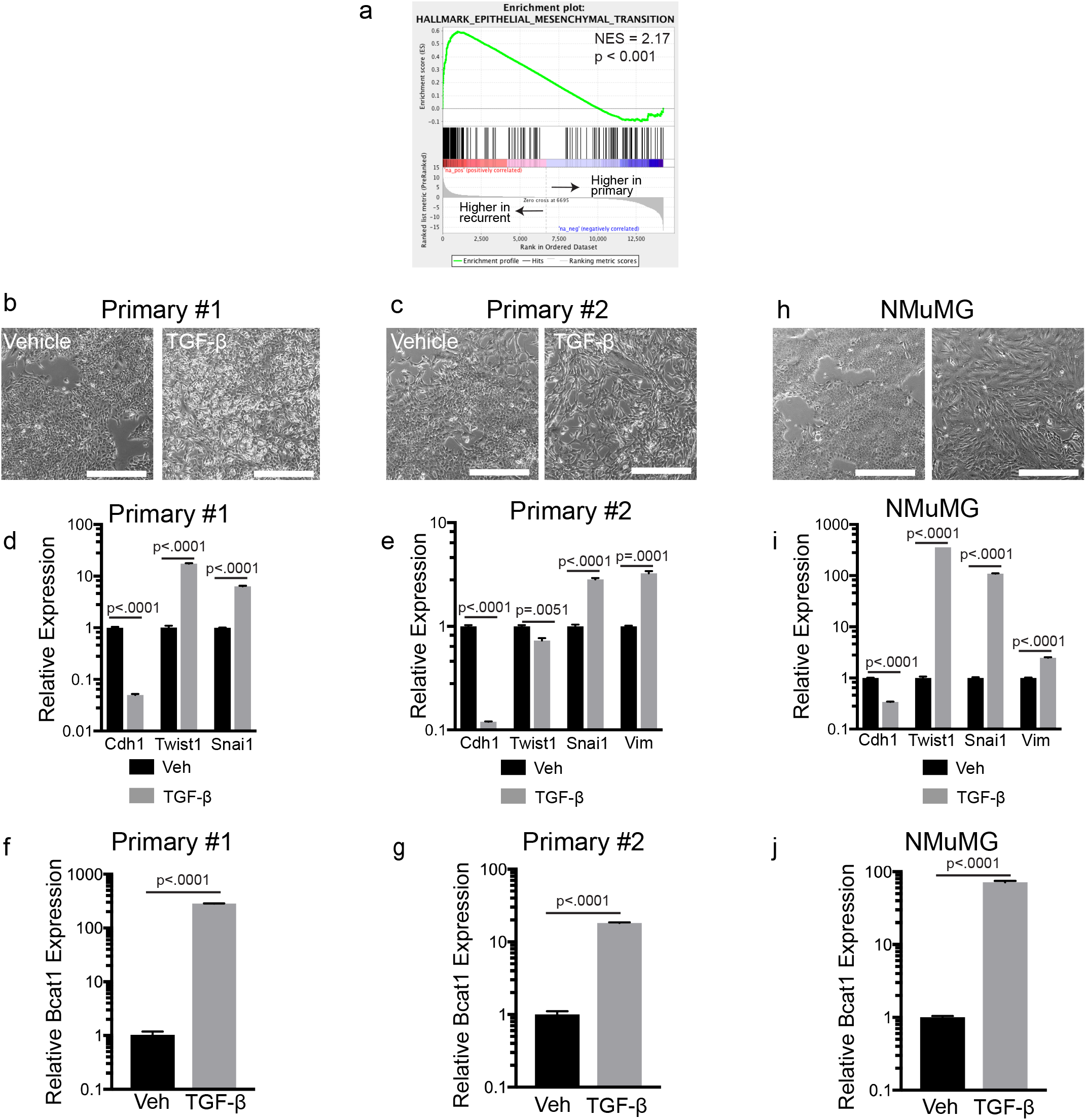
EMT induces Bcat1 upregulation. **a**, Gene set enrichment analysis of genes upregulated in recurrent tumors shows enrichment of the “Epithelial-to-Mesenchymal” gene signature in recurrent tumor cells. Enrichment scores were calculated using the Kolmogorov-Smirnov statistic and p-values were calculated using permutation testing with 1000 permutations. **b**,**c**, Brightfield microscopy images showing morphological changes induced by TGF-β treatment in two independent primary tumor cell lines. Scale bar represents 500μm. **d**,**e**, qRT– PCR analysis of expression of EMT marker genes in two independent primary tumor cell lines treated with vehicle (Veh) or TGF-β. Significance was determined by Student’s t-test. Data are representative of 2 independent experiments. **f**,**g**, qRT–PCR analysis of expression of Bcat1 in two independent primary tumor cell lines treated with vehicle (Veh) or TGF-β. Significance was determined by Student’s t-test. Data are representative of 2 independent experiments. **h**, Brightfield microscopy images showing morphological changes induced by TGF-β treatment in NMuMG cells. Scale bar represents 500μm. **i**, qRT–PCR analysis of expression of EMT marker genes in NMuMG cells treated with vehicle (Veh) or TGF-β. Significance was determined by Student’s t-test. Data are representative of 2 independent experiments. **j**, qRT–PCR analysis of expression of Bcat1 in NMuMG cells treated with vehicle (Veh) or TGF-β. Significance was determined by Student’s t-test. Data are representative of 2 independent experiments. **b-j** Mean ± SEM for n=3 biologically independent replicates.

We next asked if EMT induced expression of Bcat1 is unique to tumor cells or if it occurs in non-transformed cells. To this end, we used the normal murine mammary gland cell line NMuMG, which is a non-transformed mammary epithelial cell line [26] widely used to study EMT [27, 28]. NMuMG cells were treated with TGF-β for 10 days, and this induced morphological changes characteristic of EMT (**Fig. 3h**). qRT-PCR analysis of these cells showed that TGF-β induced downregulation of Cdh1 and upregulation of Twist1, Snail1, and Vim (**Fig. 3i**), confirming EMT transcriptional changes. Bcat1 was upregulated 72-fold relative to vehicle treated cells, demonstrating that EMT also regulates Bcat1 in non-transformed cells (**Fig. 3j**). Taken together, these results demonstrate that Bcat1 is upregulated as part of the epithelial-to-mesenchymal transition in both normal and tumor cells.

### Bcat1 expression is associated with epithelial-to-mesenchymal transition in human tumors and pan-cancer cell lines

We next wanted to substantiate our observations from mouse models using data from human tumors and cell lines. Using the RNA-seq dataset of 86 human breast cancer cell lines [22], genes were ranked based on their correlation with BCAT1 expression. Then, gene set enrichment analysis was used to identify gene signatures that correlated with BCAT1 expression. The Hallmark gene signature “epithelial-to-mesenchymal transition” was strongly correlated with BCAT1 expression (**Fig. 4a**), demonstrating that BCAT1 expression is correlated with mesenchymal markers. BCAT1 expression was anticorrelated with the Hallmark gene signatures “estrogen response early” and “estrogen response late” (**Extended Data Fig. 2a**), in agreement with the finding that BCAT1 is expressed in estrogen receptor-negative breast cancer [15]. We repeated this analysis using transcriptomic data of human breast tumors from two independent Cancer Genome Atlas (TCGA) studies [29, 30]. These analyses again clearly demonstrated a potent correlation between BCAT1 and EMT (**Fig. 4b,c**). Together, these results establish that BCAT1 expression is associated with EMT in human breast tumors.

**Fig. 4.**
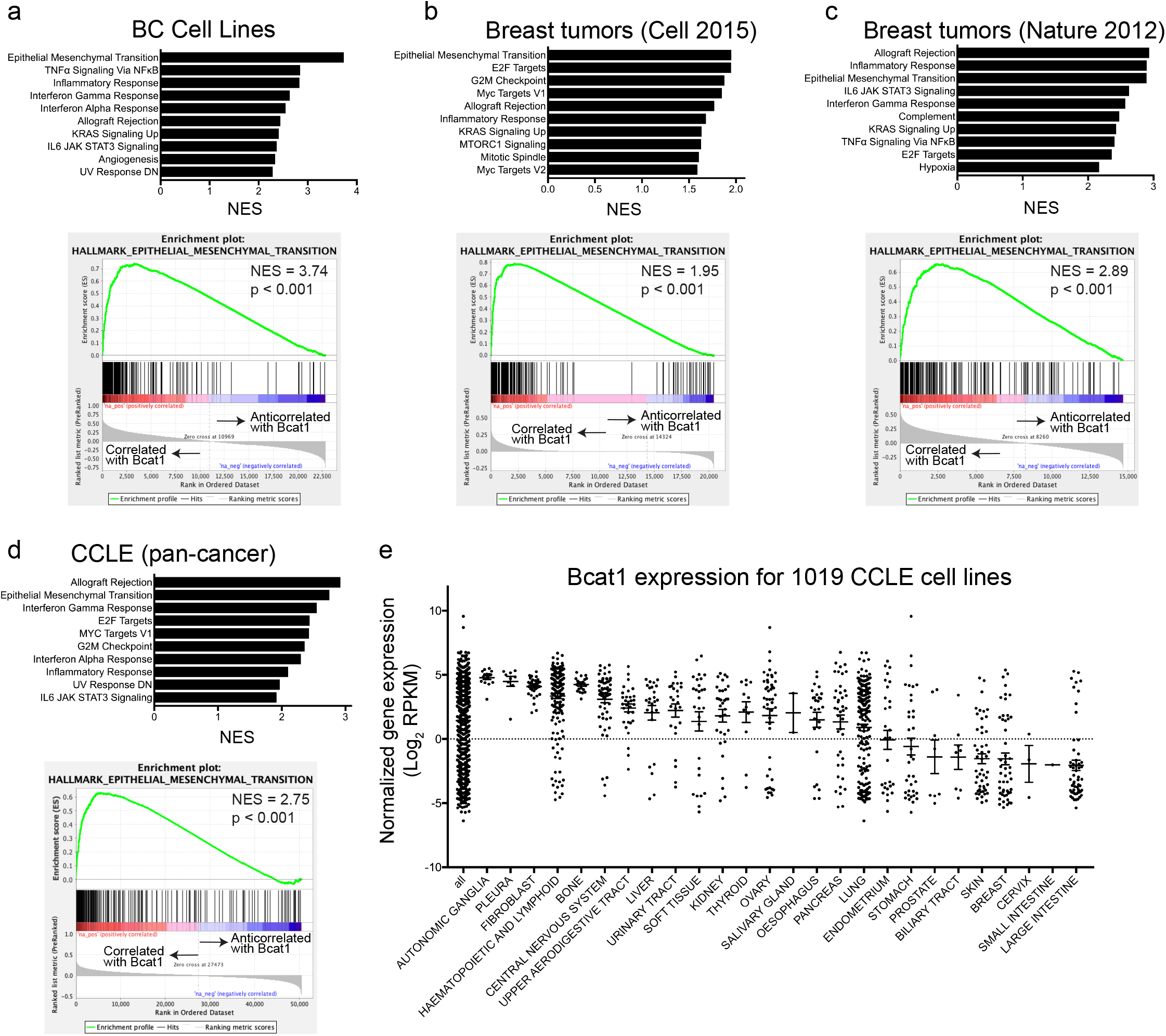
BCAT1 expression is associated with the EMT gene signature. **a**-**d**, Enrichment scores for gene signatures (top) and enrichment plots for the “Epithelial-to-Mesenchymal” gene signature (bottom) from gene set enrichment analysis of BCAT1 gene expression correlations across a panel of 86 breast cancer cell lines (**a**), breast tumors from two independent TCGA datasets (**b,c**), and the pan-cancer Cancer Cell Line Encyclopedia (CCLE) (**d**). Enrichment scores were calculated using the Kolmogorov-Smirnov statistic and p-values were calculated using permutation testing with 1000 permutations. **e**, Plot showing BCAT1 expression levels from the CCLE annotated by tissue of origin. Each dot represents the expression value for a single cell line and red bars represent the mean.

To broaden these results to other tumor types, we examined BCAT1 expression in all cell lines in the cancer cell line encyclopedia (CCLE) RNA-seq dataset [31]. For all 1076 cancer cell lines, genes were ranked based on their correlation with BCAT1, and gene set enrichment analysis was performed to identify gene sets associated with BCAT1. This analysis showed that BCAT1 expression was highly correlated with the Hallmark “epithelial-to-mesenchymal transition” gene signature (**Fig. 4d**). Plotting BCAT1 expression for each cell line based on the tissue of origin showed that many tumor types have ubiquitously high BCAT1 expression (**Fig. 4e**). Interestingly, these largely represent tumors that arise from mesodermal cells, such as the pleura, fibroblast, and bone (**Fig. 4e**). Some other tumor types derived from epithelial tissues such as ovary, lung, and pancreas, displayed very broad or bimodal distributions for BCAT1 expression, which might indicate subtype-dependent or EMT-driven regulation of BCAT1, as found in breast cancer. Taken together these results indicate that tumors that arise from mesenchymal tissues or undergo EMT during tumor progression might share a common BCAT1 dependence and be sensitive to BCAT1 inhibition.

Finally, because BCAT1 can be regulated by hypoxia in glioblastoma cells [13], we wanted to determine if hypoxia is associated with BCAT1 activation in breast cancer. Gene set enrichment analysis showed that the Hallmark “hypoxia” gene signature was not enriched in Bcat1 expressing recurrent tumor cells from MTB;TAN mice (**Extended Data Fig. 2b)**. However, the “hypoxia” gene signature was correlated with BCAT1 expression in the pan-cancer CCLE dataset (**Extended Data Fig. 2c**), although the associated was modest as compared to the “epithelial-to-mesenchymal transition” and other signatures (**Fig. 4d**). These results suggest that while hypoxia may fine-tune BCAT1 expression, EMT is responsible for the dramatic “on/off” regulation of BCAT1.

## Discussion

Here, we have interrogated the mechanism by which BCAT1 is transcriptionally activated during breast cancer recurrence. We found that it is epigenetically silenced in primary epithelial tumors driven by Her2, but recurrent tumor cells display profound epigenetic activation of BCAT1, indicating it is regulated by potent transcriptional reprogramming. Tumor recurrence in MTB;TAN mice is mediated, in part, by EMT-driven transcriptional reprogramming [18], and BCAT1 has recently been shown to be regulated by TGF-β signaling in mesenchymal fibroblast cells [25]. Interestingly, we found that TGF-β signaling was sufficient to robustly activate Bcat1 expression in primary Her2-driven tumor cells. Additionally, we found a strong correlation between BCAT1 and mesenchymal genes in human breast tumor samples, supporting the notion that EMT is the predominant mechanism by which BCAT1 expression is regulated in breast cancer cells. BCAT1 has also been reported to be transcriptionally regulated by c-MYC [10–12] and HIF [13] in carcinomas. However, c-MYC and HIF are both broadly active in many tumor type, whereas BCAT1 expression is only observed in a subset of tumors (**Figure 4e**). This suggests that c-MYC and HIF are likely to fine-tune BCAT1 expression, while EMT is responsible for its epigenetic activation.

These results suggest that primary tumors that arise from epithelial tissues that do not express BCAT1 are not likely to be dependent on BCAT1 expression if they remain in the epithelial state. However, tumor progression is often accompanied by EMT – EMT can be induced by therapy, promotes therapeutic resistance [32], and is a common mechanism driving metastatic dissemination [33]. Thus, tumor cells that undergo EMT during disease progression and tumor recurrence are likely to gain a collateral dependence on BCAT1. Pan-cancer analysis additionally demonstrated that BCAT1 expression is most pronounced in tumor types of mesenchymal tissues. It has already been reported that some of these tumor types, including lymphoma [7, 8] and glioblastoma [34] are dependent on BCAT1, and our findings that BCAT1 is activated by EMT and expressed in mesenchymal tumors broadens the potential clinical relevance of BCAT1 inhibition. Indeed, targeting the mesenchymal state has gained attention as a potential therapeutic strategy [35, 36], and these results suggest that BCAT1 inhibition might represent a promising biomarker-driven therapeutic strategy.

The function of BCAT1 during EMT or in mesenchymal cells remains to be determined. Interestingly, disrupting BCAT1 impairs tumor growth despite the presence of functional BCAT2. Furthermore, EMT has recently been associated with increased expression and activity of the branched-chain α-keto acid dehydrogenase kinase (BCKDK) [37], which inhibits mitochondrial catabolism of branched-chain α-keto acids by inactivating branched-chain α-keto acid dehydrogenase (BCKDH). These results suggest that BCAA metabolism in the cytoplasm is particularly important for tumor growth, while mitochondrial catabolism of BCAAs by BCAT2 and BCKDH is dispensable. In agreement with this, reports have shown that cytoplasmic BCAT1 activity can maintain BCAA levels to promote mTOR signaling [8], or it can deplete αKG levels to impair αKG-dependent dioxygenase activity [7]. In this latter model, BCAT1 regulates epigenetic modifying enzymes, such as DNA and histone demethylases. Epigenetic changes are critical for dynamic cell state changes, including EMT, raising the possibility that BCAT1 impacts EMT plasticity or regulates the expression of mesenchymal genes. Additionally, the reversible nature of BCAT1 transamination might allow it to buffer BCAA, BCKA and αKG levels in response to changes in nutrient availability.

## Methods

### Tissue culture and reagents

Primary and recurrent tumor cell lines were generated as described previously [1, 2]. Tumor chunks were digested with EBSS (without phenol red) supplemented with collagenase (300U/mL), hyaluronidase (100U/mL), 2% FBS, gentamycin (100μg/mL), 100U/mL Pen/Strep, and doxycycline (2μg/mL) at 37°C for 4 hours. Cells were then resuspended in Dispase II (5mg/mL) and DNase I (100μg/mL) and filtered before plating. Primary tumor cells were cultured in DMEM with 10% super calf serum, 1% Penicillin/Streptomycin, and 1% L-Glutamine supplemented with 10 ng/ml EGF, 5μg/ml insulin, 1μg/ml hydrocortisone, 5μg/ml prolactin, 1μM progesterone and 2μg/ml doxycycline to maintain HER2/neu expression. Recurrent tumor cells were cultured in DMEM with 10% SCS, 1% Penicillin/Streptomycin, and 1% L-Glutamine supplemented with 10 ng/ml EGF and 5 μg/ml insulin. NMuMG cells were obtained from the Duke University Cell Culture Facility, and sub-cultured per ATCC protocols.

### Plasmids and viral transduction for Bcat1 knockout

Lenti-Cas9-Blast, a gift from Feng Zhang (Addgene plasmid #52962) [3], was used to transduce cell with Cas9. Guide RNAs targeting Bcat1 or GFP were cloned into lentiguide-puro by Gibson Assembly (Addgene #52963) as described [4] (sgBcat1 #1 – 5’-AAGAGGATCAGAAGAAGTGG; #2 – 5’-GTAGTGCAAAACAGAGGCAG; GFP – 5’-GGGCGAGGAGCTGTTCACCG). To generate lentivirus, HEK293T cells were transfected with psPAX2 and pMDG.2 packaging plasmids (gifts from Didier Trono, EPFL, Lausanne, Switzerland; Addgene plasmids 12559 and 12660) and the lentiviral expression construct. Viral supernatant was collected after 48 and 72 hours and filtered. This virus was used to transduce cells with 6μg/mL polybrene (MilliporeSigma).

### Animals

Animal care and animal experiments were performed with the approval of, and in accordance with, guidelines of the Duke University IACUC. Mice were housed under barrier conditions with 12-hour light/12-hour dark cycles. Bitransgenic MTB;TAN mice were generated and tumors were induced as previously described [2].

### Transcriptomic analysis of MTB;TAN tumor cells

We previously published RNA sequencing data comparing two primary and two recurrent tumor cell lines derived from MTB;TAN mice [5]. The log_2_ fold change between the average expression of the two primary and two recurrent cell liens was calculated for each gene, and genes were ranked from largest (upregulated in recurrent tumor cells) to smallest (downregulated in recurrent tumor cells) before plotting or using for gene set enrichment analysis.

### qRT-PCR

RNA was extracted using the RNeasy kit (Qiagen). cDNA was synthesized using the ImProm-II Reverse Transcription System (Promega). Gene expression master mix (Applied Biosystems 4369016) and Taqman probes (ThermoFisher 4331182, Bcat1 Mm00500289_m1, Cdh1 Mm01247357_m1, Twist1 Mm00442036_m1, Snai1 Mm00441533_g1, Vim Mm01333430_m1, Actb Mm02619580_g1, Gapdh Mm99999915_g1) were used for qPCR. qPCR was performed using a CFX384 Touch Real-Time PCR Detection System (Bio-Rad). Actb and Gapdh were used for normalization controls.

### Western blotting

Western blotting was performed as previously described [2] using rabbit derived Bcat1 specific (Abcam ab110761) and a mouse derived α-Tubulin specific (Cell Signaling 3873) antibodies. Fluorophore-conjugated secondary antibodies (Life Technologies A21076 and Li-Cor 926-32210) were used for imaging using Odyssey infrared imaging system (Li-Cor).

### Cell proliferation assay

One-million recurrent tumor cells expressing Cas9 and either GFP or Bcat1 targeting sgRNAs were plated on 10cm plates. The next day, 2 plates were used for cell counting and used as the number of cells on day 0. For each timepoint, 48 hours and 120 hours, 2 plates were used to count cells, and the number of population doublings relative to day 0 was calculated.

### Orthotopic recurrent tumor growth assay

Cohorts of 14 6-week old immunocompromised mice (nu/nu) were injected orthotopically in the 4^th^ inguinal mammary fat pad with 1×10^5^ recurrent tumor cells expressing Cas9 and either GFP or Bcat1 targeting sgRNA. Tumor growth was monitored by palpation. Tumor AUC was calculated using the formula [(vol_1_ + vol_2_)/2]*(day_2_-day_1_).

### EMT induction

Primary tumor cells and NMuMG cells were treated with 5ng/mL mouse TGF-β1 (Cell Signaling Technologies # 5231) or vehicle (20 mM citrate, pH 3.0) for 10 days. Morphological changes were confirmed by brightfield microscopy, and cells were then plated for RNA isolation and protein isolation.

### Analysis of Bcat1 expression in cancer cell lines

For analysis of breast cancer cell lines, publicly available RNA-sequencing data from a panel of 86 breast cancer cell lines [6] were downloaded from http://neellab.github.io/bfg/, and the log_2_ expression value was plotted for BCAT1 and other metabolic genes. Minimum value was set at log_2_(0.1) FPKM by Marcotte et al. [6]. For pan-cancer analysis, publicly available RNA-sequencing data from a panel of 1072 pan-cancer cell lines [7] were downloaded from the Broad Institute Cancer Cell Line Encyclopedia https://portals.broadinstitute.org/ccle/data, and the log_2_ expression value was plotted for BCAT1 and other metabolic genes.

### ChIP-Seq Analysis of MTB;TAN tumor cell lines

We previously published ChIP-sequencing data for primary and recurrent tumor cell lines. These ChIP-seq data were processed as described [5]. Briefly, fastq files were processed with TrimGalore [8], Bowtie [9], and the MACS2 peak-calling algorithm [10]. Bam files were converted into Bigwig format by binning reads into 100 bp segments. Images were generated in the IGV desktop viewer (Broad Institute). ChIP sequencing data are available online using the NCBI Short Read Archive (SRA) under project accession number PRJNA505839.

### Bcat1 correlation analysis

Publicly available gene expression data sets for 86 breast cancer cell lines [6] and 2 independent human breast tumor studies [11, 12] were used. For each dataset, genes were ranked based on their correlation (Pearson) with BCAT1 expression, and this ranked list was used for gene set enrichment analysis to identify gene signatures that are associated with BCAT1 expression.

### Statistics and Reproducibility

Plots and statistical analyses were generated using Prism 7 software. Specific statistical tests are identified in figure legends for each experiment. For all western blots and qRT-PCR experiments, a single experiment is shown that is representative of results from 3 independent experiments, unless otherwise noted in the figure legend.

## Data Availability

RNA- and ChIP sequencing data comparing primary and recurrent MTB;TAN-derived tumor cell lines is available online using the NCBI Short Read Archive (SRA) under project accession number PRJNA505839 [5]. RNA sequencing data comparing gene expression for 86 breast cancer cell lines are available from http://neellab.github.io/bfg/ [6]. RNA sequencing data for 1072 pan-cancer cell line analysis were downloaded from the Broad Institute Cancer Cell Line Encyclopedia at https://portals.broadinstitute.org/ccle [7].

## Acknowledgements

We thank S. Y. Kim from the Duke Functional Genomics Core for designing and cloning sgRNA. We thank N. Mabe for his advice and bioinformatic analyses. This work was funded by National Cancer Institute grants F31CA228321 (D.B.F.) and R01CA208042 (J.V.A.), and the V Foundation, American Cancer Society (132556-RSG-18-130-CCG), the Integrative metabolomics shared resource, and by startup funds from the Duke Cancer Institute, the Duke University School of Medicine and the Whitehead Foundation (to J.V.A.).

## Author Contributions

J.V.A. and D.B.F. were responsible for the conception, design and interpretation of all experiments. D.B.F. performed all experiments and collected data. D.B.F. and J.V.A. wrote the manuscript. J.V.A. supervised all work.

## Extended Data Figure Legends

**Extended Data Fig. 1.**
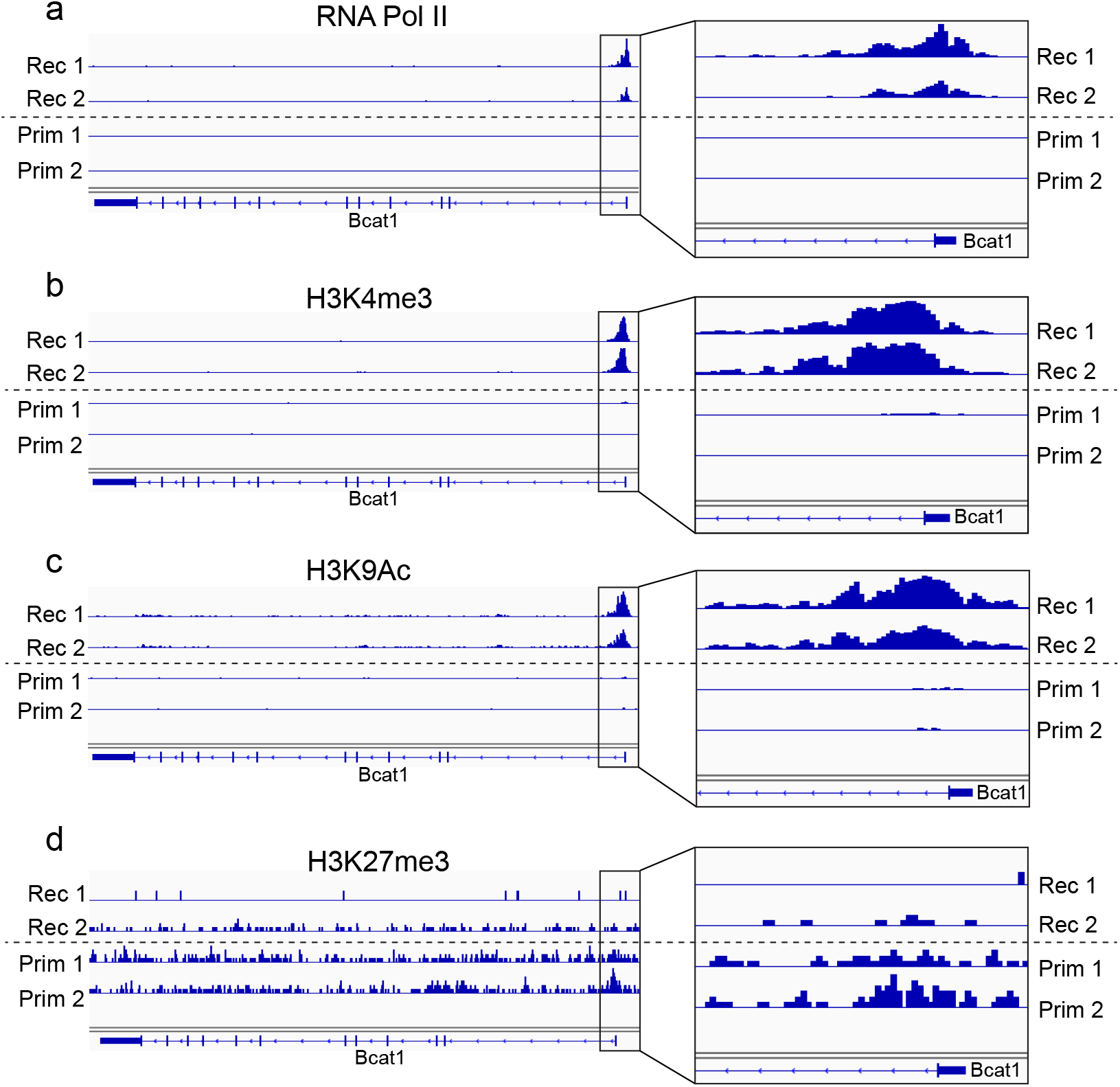
Dynamic epigenetic changes regulate Bcat1. **a-d**, ChIP-seq plots showing RNA Pol II occupancy (**a**), H3K4me3 abundance (**b**), H3K9Ac abundance (**c**), and H3K27me3 abundance (**d**) for the Bcat1 gene (left) and promoter region (right) in two recurrent and two primary MTB;TAN cell lines.

**Extended Data Fig. 2.**
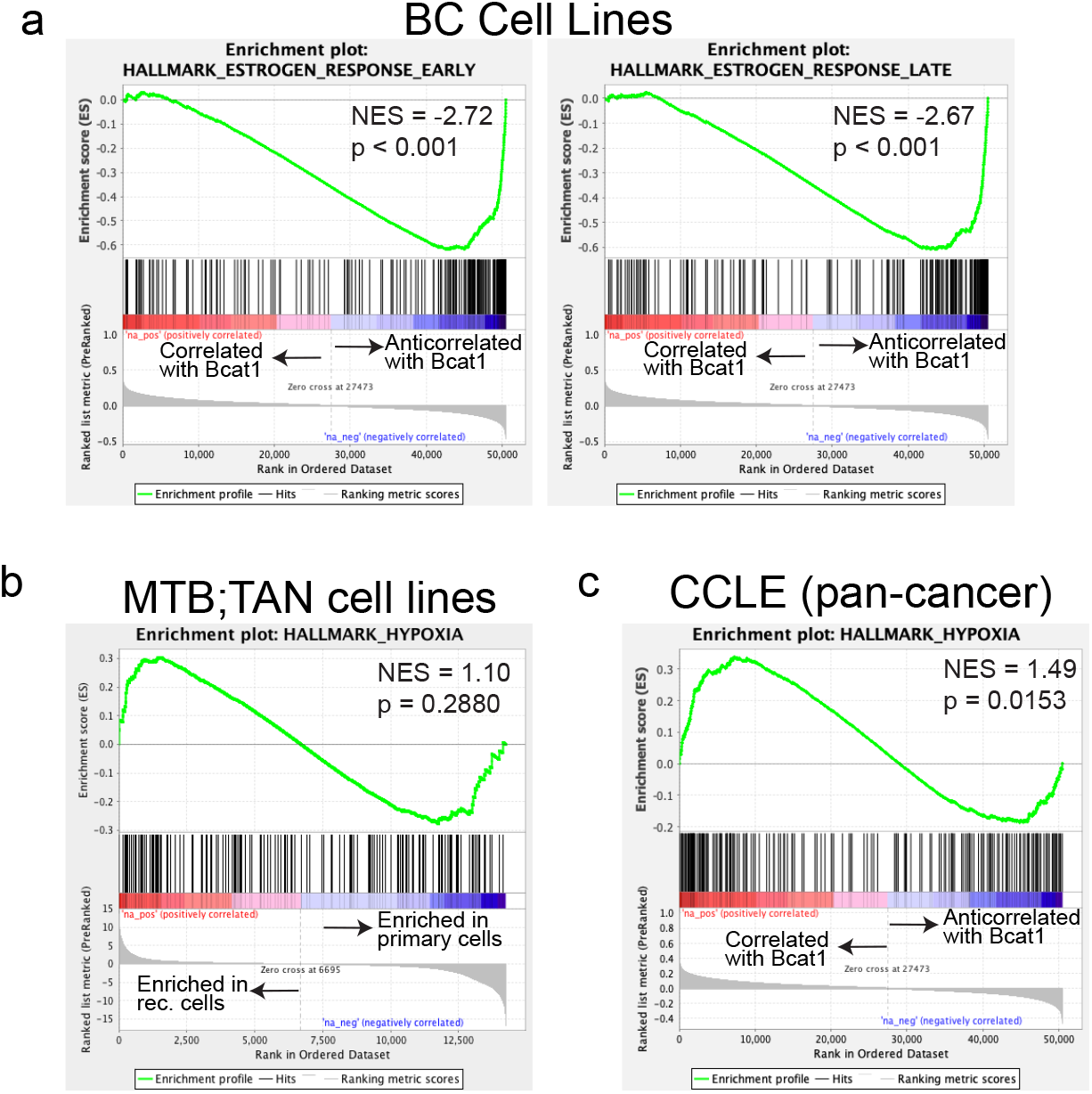
BCAT1 expression is not robustly correlated with hypoxia. **a**, Enrichment plots for the “Estrogen Response Early” and “Estrogen Response Late” gene signatures from gene set enrichment analysis of BCAT1 gene expression correlations for a panel of 86 breast cancer cell lines. **b**, Enrichment plot for the “Hypoxia” gene signature from gene set enrichment analysis comparing primary and recurrent MTB/TAN cell lines. **c**, Enrichment plot for the “Hypoxia” gene signature from gene set enrichment analysis of BCAT1 gene expression correlations for the pan-cancer Cancer Cell Line Encyclopedia (CCLE) database. **a-c**, Enrichment scores were calculated using the Kolmogorov-Smirnov statistic and p-values were calculated using permutation testing with 1000 permutations.

